# Type I interferon signaling induces a delayed antiproliferative response in Calu-3 cells during SARS-CoV-2 infection

**DOI:** 10.1101/2023.02.28.530557

**Authors:** Juliana Bragazzi Cunha, Kyle Leix, Emily J. Sherman, Carmen Mirabelli, Andrew A. Kennedy, Adam S. Lauring, Andrew W. Tai, Christiane E. Wobus, Brian T. Emmer

## Abstract

Disease progression during SARS-CoV-2 infection is tightly linked to the fate of lung epithelial cells, with severe cases of COVID-19 characterized by direct injury of the alveolar epithelium and an impairment in its regeneration from progenitor cells. The molecular pathways that govern respiratory epithelial cell death and proliferation during SARS-CoV-2 infection, however, remain poorly understood. We now report a high-throughput CRISPR screen for host genetic modifiers of the survival and proliferation of SARS-CoV-2-infected Calu-3 respiratory epithelial cells. The top 4 genes identified in our screen encode components of the same type I interferon signaling complex – *IFNAR1*, *IFNAR2*, *JAK1*, and *TYK2*. The 5^th^ gene, *ACE2*, was an expected control encoding the SARS-CoV-2 viral receptor. Surprisingly, despite the antiviral properties of IFN-I signaling, its disruption in our screen was associated with an increase in Calu-3 cell fitness. We validated this effect and found that IFN-I signaling did not sensitize SARS-CoV-2-infected cultures to cell death but rather inhibited the proliferation of surviving cells after the early peak of viral replication and cytopathic effect. We also found that IFN-I signaling alone, in the absence of viral infection, was sufficient to induce this delayed antiproliferative response. Together, these findings highlight a cell autonomous antiproliferative response by respiratory epithelial cells to persistent IFN-I signaling during SARS-CoV-2 infection. This response may contribute to the deficient alveolar regeneration that has been associated with COVID-19 lung injury and represents a promising area for host-targeted therapeutic development.

## INTRODUCTION

The respiratory epithelium plays an important role during COVID-19, as alveolar epithelial cells express the SARS-CoV-2 receptor ACE2 and support active viral replication during infection^1, 2^. Diffuse alveolar damage, characterized by the injury of epithelial cells, compromised barrier function, and leakage of proteinaceous exudate into the alveolar space, is a histopathologic hallmark of severe COVID-19 and other causes of the acute respiratory distress syndrome (ARDS)^1, 3^. Injury to the alveolar epithelium is counterbalanced by a regenerative pathway involving the proliferation and trans-differentiation of type 2 alveolar epithelial (AT2) cells that repopulate the alveolar lining and resorb fluid from the alveolar space^4–7^. COVID-19 has been associated with a deficiency in this AT2 regenerative response^8, 9^, though the molecular basis for this deficiency is unknown.

Severe cases of COVID-19 typically develop 7 or more days after initial infection^10^, at which time an exuberant host response, rather than uncontrolled viral replication, appears to drive lung pathology. Accordingly, virus-targeted therapies such as remdesivir have little or no benefit in severe COVID-19^11, 12^, while host-targeted therapies such as corticosteroids and Janus kinase inhibitors have been shown to improve clinical outcomes^13–16^. These treatments modulate several host signaling pathways, however, and it is unknown which are responsible for their therapeutic effect. The type I interferon (IFN-I) response has been widely studied, but its precise role in SARS-CoV-2 infection remains enigmatic and likely depends on the timing and magnitude of the response^17^.

Over the past decade, CRISPR screening has emerged as a powerful tool for detecting functional interactions between host cells and viral pathogens^18^. Since the onset of the COVID-19 global pandemic, several groups have applied this approach to SARS-CoV-2 infection^19–29^. Although some genes have been consistently identified across multiple studies, many have shown limited overlap, potentially due to differences in cell type or other variables in screen design. For most identified host factors, their mechanism of effect during viral infection remains unclear.

We now report the results of a CRISPR screen for host factors affecting SARS-CoV-2 infection in Calu-3 cells, a cell line of respiratory epithelial origin that recapitulates the SARS-CoV-2 replication kinetics and interferon responses of primary human airway cells^30^. The findings from our screen and subsequent mechanistic studies implicate IFN-I signaling as both necessary and sufficient for a delayed antiproliferative response that limits the recovery of Calu-3 cells that survive the initial peak of viral replication and cell death.

## RESULTS

### CRISPR screen for modifiers of Calu-3 cell fitness during SARS-CoV-2 infection

We previously synthesized a high-resolution CRISPR library targeting 833 genes implicated in SARS-CoV-2 biology^31^ based on their identification in prior CRISPR screens of SARS-CoV-2 cytopathic effect^19, 20, 22–26, 32^, human genome-wide association studies of COVID-19 susceptibility^33^, RNA-seq analysis of genes whose expression correlated with the SARS-CoV-2 receptor ACE2^34^, and our own genome-wide CRISPR screen for modifiers of ACE2 abundance^31^. Our earlier application of this library to a flow cytometry-based screen led to our successful identification of cell type-specific modifiers of ACE2 surface abundance in HuH7 and Calu-3 cells^31^. In this study, we repurposed this same library to screen for modifiers of Calu-3 cell fitness during SARS-CoV-2 infection (Fig 1A). We generated pools of CRISPR-edited cells that we then infected with the WA1 strain of SARS-CoV-2 at an MOI of 1. SARS-CoV-2-infected cells began exhibiting signs of cytopathic effect at day 2 that became most prominent at approximately day 4 post-infection. Surviving cells were maintained in culture for 14 days post-infection to allow for outgrowth of edited cells with a fitness advantage. We then quantified the abundance of each individual gRNA by massively parallel sequencing from genomic DNA isolated at days 0 and 14 post-infection and analyzed gRNA enrichment using MAGeCK^35^ (Fig 1B, Supplemental Tables 1-2). Quality control analysis confirmed adequate depth of sampling (Fig S1A) with high reproducibility between independent biologic replicates (Fig 1C) and robust discrimination of the positive control *ACE2*-targeting and negative control nontargeting gRNAs in SARS-CoV-2 infected cells (Fig 1D). To avoid identifying gene perturbations that had a general influence on host cell fitness independently of SARS-CoV-2 infection, we filtered out genes targeted by gRNAs that were significantly enriched or depleted in day 0 samples relative to the CRISPR library plasmid pool (Fig S2A, Supplemental Tables 3-4). The genes removed by this filtering strategy showed considerable overlap with those identified as serving a core essential function in an analysis of 1070 cell lines^36^ (Fig S2B).

**Figure 1.**
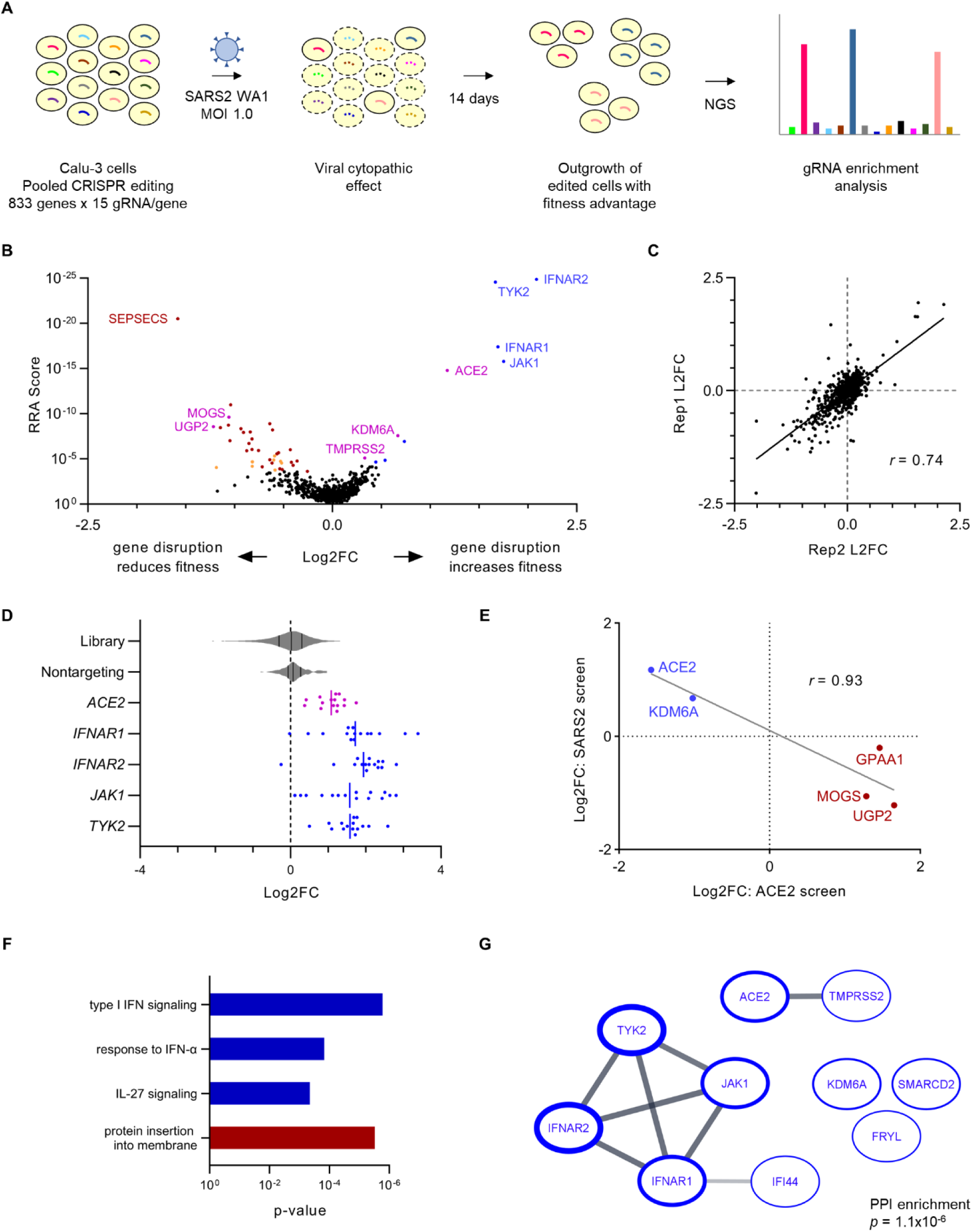
CRISPR screen for modifiers of Calu-3 cell fitness during SARS-CoV-2 infection. (A) Schematic of CRISPR screening strategy. (B) Volcano plot of CRISPR screen results. Genes whose disruption confers a significant (FDR<5%) increase or decrease in cell fitness during SARS-CoV-2 infection are highlighted in blue or red, respectively. Filtered genes whose disruption confers a significant effect on cell fitness in the absence of viral infection (Fig S2) are shaded in orange. Identified genes with a known role in SARS-CoV-2 entry or ACE2 regulation are highlighted in magenta. Select genes are labeled and full screen results are provided in Supplemental Tables 1 and 2. (C) Pairwise correlation between independent biologic replicates for the aggregate log2 fold-change of all gRNAs targeting a given gene. Correlation coefficient was calculated by Pearson method. (D) Individual gRNA log2 fold-change for each of 15 gRNAs targeting the indicated gene, with violin plots showing the distribution of changes for the entire library and the subset of control nontargeting gRNAs. Vertical lines indicate mean of individual gRNAs for each gene. (E) Correlation between aggregate gRNA log2 fold-change in independent screens for modifiers of Calu-3 ACE2 surface abundance and cell fitness during SARS-CoV-2 infection. Genes whose disruption was associated with a decrease in surface ACE2 and an increase in cell fitness during SARS-CoV-2 infection are highlighted in blue. Genes whose disruption was associated with an increase in surface ACE2 and a decrease in cell fitness during SARS-CoV-2 infection are highlighted in red. Correlation coefficient was calculated by Pearson method. (F) Functional annotations significantly enriched (p < 10^-3^) among identified genes whose disruption was associated with a significant increase (blue) or decrease (red) in Calu-3 cell fitness during SARS-CoV-2 infection. For multiple nodes within the same hierarchy, the annotation with the most significant enrichment was selected. (G) Established protein-protein interactions in the STRING database among genes identified in the screen whose disruption was associated with a significant increase in Calu-3 cell fitness during SARS-CoV-2 infection. Borders of individual nodes are weighted by the -log(RRA score) in the screen, and lines connecting nodes are weighted by the strength of the protein-protein interaction within the STRING database. The significance of the number of protein-protein interactions relative to a randomly selected gene set was calculated by STRING.

Overall, we identified 10 genes whose CRISPR targeting was associated with a significant (FDR<5%) increase in Calu-3 cell fitness and 31 genes whose targeting was associated with a significant decrease in fitness during SARS-CoV-2 infection (Fig 1B, Supplemental Tables 1-2). As expected, top hits of the former group included both *ACE2* and *TMPRSS2*, two genes with well-characterized roles in SARS-CoV-2 entry^37^. Similarly, among the genes we previously identified as modifiers of ACE2 surface abundance in Calu-3 cells^31^, we detected a strong inverse correlation between their effect on ACE2 and their effect on cell fitness during SARS-CoV-2 infection (Fig 1E). This correlation supports both the validity of the screen and the relevance of quantitative changes in ACE2 abundance to cellular sensitivity to SARS-CoV-2 cytopathic effect.

In addition to *ACE2* and the ACE2 modifiers discussed above, we identified several other genes that influenced the fitness of SARS-CoV-2-infected Calu-3 cells. Ontology analysis revealed a significant enrichment for a limited number of functional annotations, with the greatest effect observed for the type I interferon signaling pathway (Fig 1F). Analysis of the STRING database likewise revealed a significant enrichment in established protein-protein interactions (Fig 1G, Fig S3), most prominently among components of the type I interferon receptor signaling complex encoded by *IFNAR1*, *IFNAR2*, *JAK1*, and *TYK2*. Unexpectedly and in contrast to the canonical antiviral properties of the type I interferon response, disruption of type I interferon signaling was associated with an increase in Calu-3 fitness, with gRNAs targeting these genes consistently becoming enriched during SARS-CoV-2 infection (Fig 1D). The aggregate magnitude of enrichment at 14 days post-infection for gRNAs targeting each of these genes was even greater than that observed for *ACE2* (Fig 1D). These findings indicate that despite any potential antiviral effects of endogenous IFN-I signaling, its overall impact on SARS-CoV-2-infected populations in our screen led to a decrease in Calu-3 cell fitness.

### The overall effect of IFN-I signaling on SARS-CoV-2-infected Calu-3 cells is context-dependent

To validate our screen finding that IFN-I signaling decreased Calu-3 cell fitness during SARS-CoV-2 infection and to explore the mechanism for this effect, we next targeted the endogenous *IFNAR1* locus of Calu-3 cells by CRISPR and confirmed the efficient depletion of *IFNAR1* mRNA (Fig 2A) (consistent with nonsense-mediated decay of transcripts with frameshift-causing indels) and IFNAR1 protein (Fig 2B). We also confirmed that *IFNAR1*-disrupted cells were resistant to the induction of interferon-stimulated gene expression by exogenous β-IFN at a concentration approximating the endogenous response of SARS-CoV-2-infected wild-type Calu-3 cells^30^ (Fig 2A, 2B). Paradoxically, despite the protective effect of *IFNAR1* disruption in our screen, testing of these *IFNAR1*-disrupted cells in parallel to control cells revealed a greater decrease in viable cells during SARS-CoV-2 infection (Fig 2C). A similar sensitization to SARS-CoV-2 cytopathic effect was observed for pharmacologic pre-treatment with a Janus kinase inhibitor, baricitinib, that inhibited IFN-I signaling in Calu-3 cells (Fig 2D, 2E).

**Figure 2.**
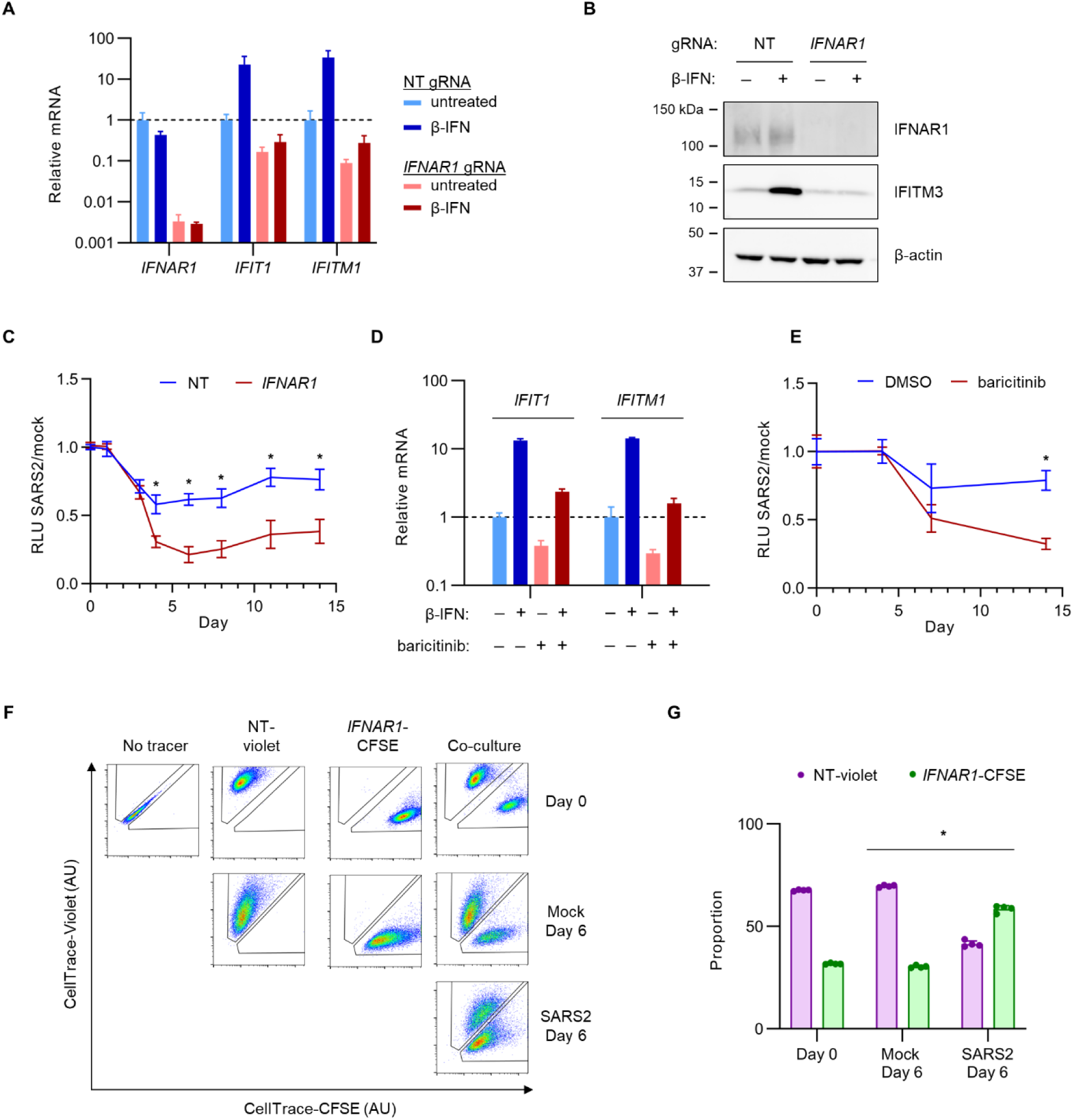
*IFNAR1*-disrupted Calu-3 cells exhibit reduced fitness during SARS-CoV-2 infection when cultured separately from control cells but increased fitness when cultured together. (A-B) Calu-3 cells were transduced with lentiviral constructs delivering Cas9 and either a *IFNAR1*-targeting or nontargeting control gRNA, followed by treatment with 1000 U/mL exogenous β-IFN or vehicle control for 48 hr and analysis of (A) mRNA by qRT-PCR with primer pairs for the indicated gene and (B) protein lysates by immunoblotting with antibodies for the indicated proteins. (C) Control and *IFNAR1*-disrupted cells were seeded in 96 well plates, infected with SARS-CoV-2 at MOI of 1 or mock-infected, and the resulting number of viable cells at indicated time points assayed by CellTiter-Glo luminescence. (D) Calu-3 cells were treated with 1000 U/mL β-IFN or vehicle in the presence of 1 µM baricitinib or vehicle for 48 hr and tested for induction of interferon-stimulated gene mRNA expression qRT-PCR. (E) Calu-3 cells were treated with1 µM baricitinib or vehicle at the onset of infection with SARS-CoV-2 WA1 strain at MOI of 1. The number of viable cells at the indicated time points was monitored by CellTiter-Glo luminescence. (F-G) Calu-3 cells were treated with a nontargeting gRNA or *IFNAR1*-targeting gRNA, loaded with CellTrace-Violet or CellTrace-CFSE, respectively, and either maintained in isolation as gating controls or pooled together. The relative proportion of each cell type in the mixture was visualized (F) and quantified (G) by flow cytometry. Error bars represent standard deviations of 2-4 replicates for each assay and asterisks indicate p < 0.001 by Student’s t-test.

The discrepancy between our screen findings and our subsequent validation studies led us to consider the differences in experimental design between these approaches. In the screen, *IFNAR1*-disrupted cells shared the same extracellular environment as neighboring cells with intact IFN-I signaling, whereas in single gene disruption studies, each population occupied a separate physical space with its own conditioned media. To test whether this difference might underlie the discordance in our findings, we next tested control and *IFNAR1*-disrupted cells in a competitive co-culture experiment. We loaded each cell type with a different fluorescent dye and tracked their relative proportions over the course of SARS-CoV-2 or mock infection by flow cytometry. Consistent with our screen and in contrast to the observations with separate cultures, we found that *IFNAR1*-disrupted cells in this context exhibited a clear competitive advantage over control cells during SARS-CoV-2 infection but not during mock infection (Fig 2F-G, S4). This competitive advantage for *IFNAR1*-disrupted cells was not mediated by the fluorescent dyes themselves, as it was observed with either combination of cell population and dye (Fig S5). Together, these findings validate the detection of IFN-I signaling genes as modifiers of Calu-3 cell fitness in our pooled screen but indicate that the protective effect of IFN-I signaling disruption is dependent on the presence of neighboring cells with intact IFN-I signaling.

### The early IFN-I response protects SARS-CoV-2-infected Calu-3 cells by restricting viral replication

The modifying influence of a separate or shared environment for *IFNAR1*-disrupted cells with control cells suggested a complexity of interactions with the extracellular space during SARS-CoV-2 infection. We reasoned that if *IFNAR1* disruption led to both a fitness-promoting cell autonomous signal and a fitness-reducing paracrine signal, then only the former effect would be isolated in a co-culture experiment in which both cell types shared the same media. When cultured separately, however, the impact on cell fitness would reflect the net effect of both signals. We therefore tested how endogenous *IFNAR1* disruption affected the extracellular space during SARS-CoV-2 infection. We found that conditioned media from *IFNAR1*-disrupted cells contained greater numbers of infectious virions, with both a higher peak at day 2 post-infection and a longer persistence at subsequent time points (Fig 3A). Consistent with these findings, we also observed for *IFNAR1*-disrupted cells an increase in SARS-CoV-2 RNA associated with both the conditioned media (Fig 3B) and cell monolayer (Fig 3C), as well as an increase in luminescence during infection with a SARS-CoV-2 nanoluciferase reporter virus^38^ (Fig 3D). These findings support the importance of the endogenous IFN-I response in restricting SARS-CoV-2 replication in Calu-3 cells, and suggest that in single cultures, any fitness-promoting effect conferred by IFN-I signaling disruption may be offset by an increase in viral replication.

**Figure 3.**
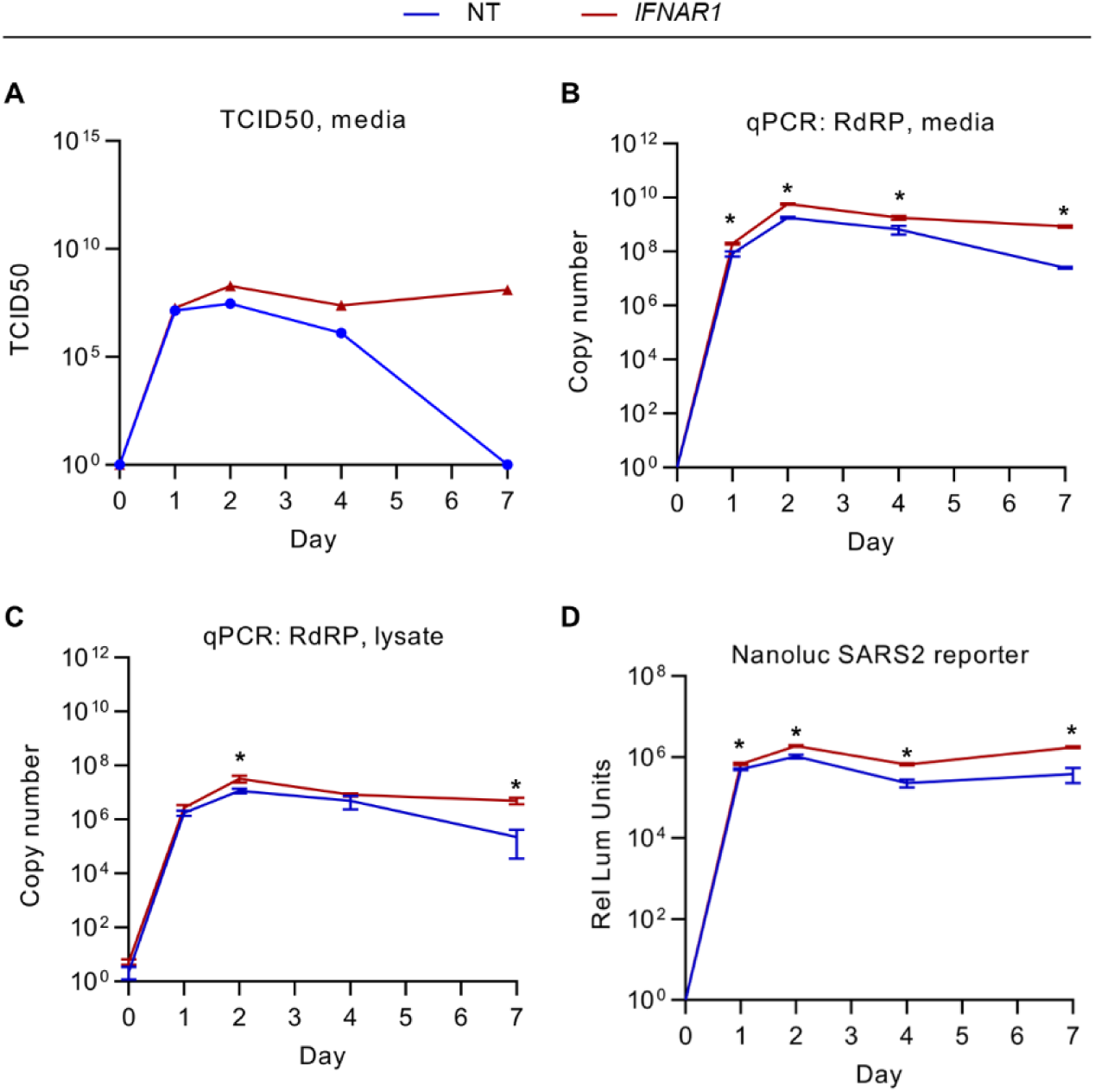
Early IFN-I signaling restricts SARS-CoV-2 replication in Calu-3 cells. (A-C) Control nontargeting and *IFNAR1*-disrupted cells were infected with SARS-CoV-2 WA strain at MOI of 1 and cellular lysates and conditioned media collected at the indicated time points and quantified for (A) the quantity of infectious virions in conditioned media by TCID50 in VeroE6 cells and (B-C) SARS-CoV-2 RdRP copy number in (B) cellular RNA and (C) RNA isolated from conditioned media. (D) Calu-3 cells treated with nontargeting or *IFNAR1*-targeting gRNAs were infected with a SARS-CoV-2 nanoluciferase reporter strain and the resulting luminescence signal assayed at various time points post-infection. (B-D) Error bars represent standard deviations of 4-6 replicates for each assay and asterisks indicate p < 0.05 by Student’s t-test.

### Persistent IFN-I signaling during SARS-CoV-2 infection limits the expansion of Calu-3 cells that survive early cytopathic effect

To better understand the basis for the fitness-decreasing effect of IFN-I signaling during SARS-CoV-2 infection, we next performed a detailed analysis of co-cultured control and *IFNAR1*-disrupted cells over the course of infection. We calculated absolute cell numbers by normalization to the event frequency of flow cytometry beads spiked into samples at a known concentration. Both control and *IFNAR1*-disrupted cells increased in number over the first 2 days of infection before reaching a plateau and then decreasing by day 4 post-infection (Fig 4A), corresponding to the visual appearance of cytopathic effect in these cells. Subsequently, the number of control cells remained low throughout the remainder of the experiment until decay in the fluorescent signal precluded analysis beyond day 8. *IFNAR1*-disrupted cells, by contrast, underwent a robust increase in cell numbers from day 4 until day 8.

**Figure 4.**
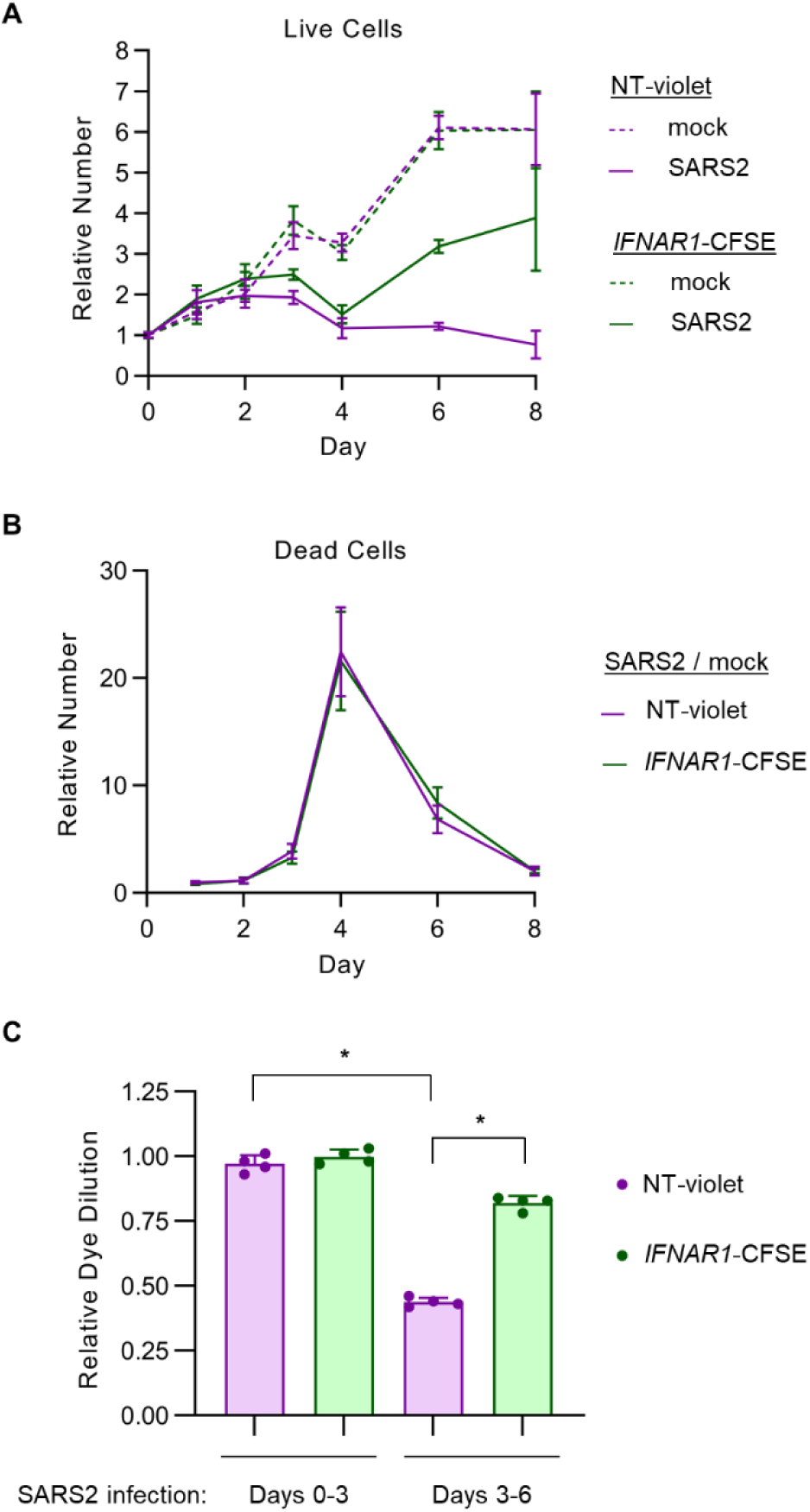
Persistent IFN-I signaling induces a delayed antiproliferative effect in SARS-CoV-2-infected Calu-3 cells. (A-B) Calu-3 cells were treated with a control nontargeting or *IFNAR1*-targeting gRNA and loaded with CellTrace-Violet or CellTrace-CFSE, respectively, pooled together, and either infected with SARS-CoV-2 WA1 strain at MOI of 1 or mock-infected. The absolute number of cells for each population within the co-culture was calculated at each time point by flow cytometry with comparison of event frequency to standard beads spiked into each sample at a known density. Live and dead cells were discriminated by staining with Live-Dead Fixable Far Red viability dye. (A) Live cells adherent in the cell monolayer were normalized for each population to the beginning quantity of viable cells at day 0. (B) Dead cells detached in the conditioned media are plotted for SARS-CoV-2-infected conditions relative to mock-infected conditions at the same time point. (C) Proliferation of each cell population within the co-coculture was quantified by the relative dilution of fluorescent dye over the indicated time interval, normalized to the proliferation of that cell type over the first 3 days of mock-infection. Error bars represent standard deviations of 4 independent infections and asterisks indicate p < 0.001 by Student’s t-test.

The delayed fitness advantage of *IFNAR1*-disrupted cells relative to control cells might have been caused by a decrease in cell death and/or an increase in cell proliferation. To distinguish between these possibilities, we tracked the absolute number of dead cells in either population over the course of SARS-CoV-2 infection. For each population, we observed far fewer dead cells associated with the cell monolayer than in the conditioned media (Fig S6), suggesting that cell death led to rapid detachment of these cells from the plate. Relative to mock-infected conditions, the number of dead cells in the media rose abruptly at day 3 and peaked at day 4 post-infection (Fig 4B), corresponding to the timing of the visual appearance of cytopathic effect and the decrease in viable cells associated with the monolayer (Fig 4A). Compared to co-cultured control cells, *IFNAR1*-disrupted cells were equally sensitive to SARS-CoV-2-associated cell death (Fig 4B).

We next monitored cellular proliferation of control and *IFNAR1*-disrupted cells over time by measuring the dilution of either fluorescent dye with each cell doubling. Both cell types exhibited comparable rates of proliferation over the first 3 days of either mock or SARS-CoV-2 infection (Fig 4C). Between days 3 and 6, however, SARS-CoV-2-infected control cells exhibited a clear decrease in dye dilution (Fig 4C), corresponding to their lack of proliferation during this time (Fig 4A). *IFNAR1*-disrupted cells, by contrast, were resistant to this decrease in dye dilution between days 3 and 6 post-infection (Fig 4C), consistent with the expansion of this population during this time (Fig 4A). Together, these findings indicate that persistent IFN-I signaling during SARS-CoV-2 infection reduces the fitness of Calu-3 cells through an antiproliferative response rather than a sensitization to cell death.

### Analysis of factors influencing the fitness advantage of *IFNAR1*-disrupted Calu-3 cells during SARS-CoV-2 infection

Recently, three other studies have reported high-throughput CRISPR screens for modifiers of Calu-3 cell fitness during SARS-CoV-2 infection^27–29^. We compared the genes identified in our screen with the findings from each of these studies (Fig S7). Although *ACE2* was a top hit in each screen, and fitness-promoting and fitness-reducing gene perturbations were broadly correlated between the studies, there was otherwise a modest overlap between the results of our screen and these studies. Like us, Biering et al detected a significant association between disruption of *IFNAR1* and *IFNAR2* with increased Calu-3 cell fitness during SARS-CoV-2 infection, though with a magnitude of enrichment less than we observed^28^. Rebendenne et al did not detect *IFNAR1*, *IFNAR2*, *JAK1*, or *TYK2* among the top hits of their primary genome-wide screen, though interestingly they did identify *IRF9*, which is downstream of canonical IFN-I signaling, as the top hit of their secondary screen despite this gene showing no significant effect in the primary genome-wide screen^29^. *IRF9* was also identified by Biering et al, along with the other genes encoding its trimeric transcription factor complex - *STAT1* and *STAT2*^28^. Israeli et al did not detect a significant effect for disruption of IFN-I signaling during SARS-CoV-2 infection of Calu-3 cells^27^. Each of these studies had technical differences in experimental design, including the multiplicity of infection, the plating density of cells, and the duration of culturing after infection before gRNA analysis.

To analyze how different experimental parameters might influence the competitive advantage of IFN-I signaling disruption during Calu-3 SARS-CoV-2 infection, we again tracked the relative abundance of co-cultured control and *IFNAR1*-disrupted cells under different conditions. We found that lower plating densities of Calu-3 cells led to a greater competitive advantage for *IFNAR1*-disrupted cells (Fig 5A), associated with a greater degree of proliferation for these cells (Fig 5B). The amount of virus used for infection also had a large modifying influence, with increasing MOI leading to a more pronounced proliferation advantage for *IFNAR1*-disrupted cells (Fig 5C-D). The competitive advantage of *IFNAR1*-disrupted cells was not unique to the WA1 strain, as it was also observed for each of the Alpha, Beta, Gamma, and Omicron variants of SARS-CoV-2, albeit with varying magnitude of effect (Fig 5E-F). These findings suggest that experimental conditions that are associated with more cell doublings after the initial cytopathic effect and greater production of IFN-I are more likely to reveal a proliferation advantage conferring by IFN-I signaling disruption.

**Figure 5.**
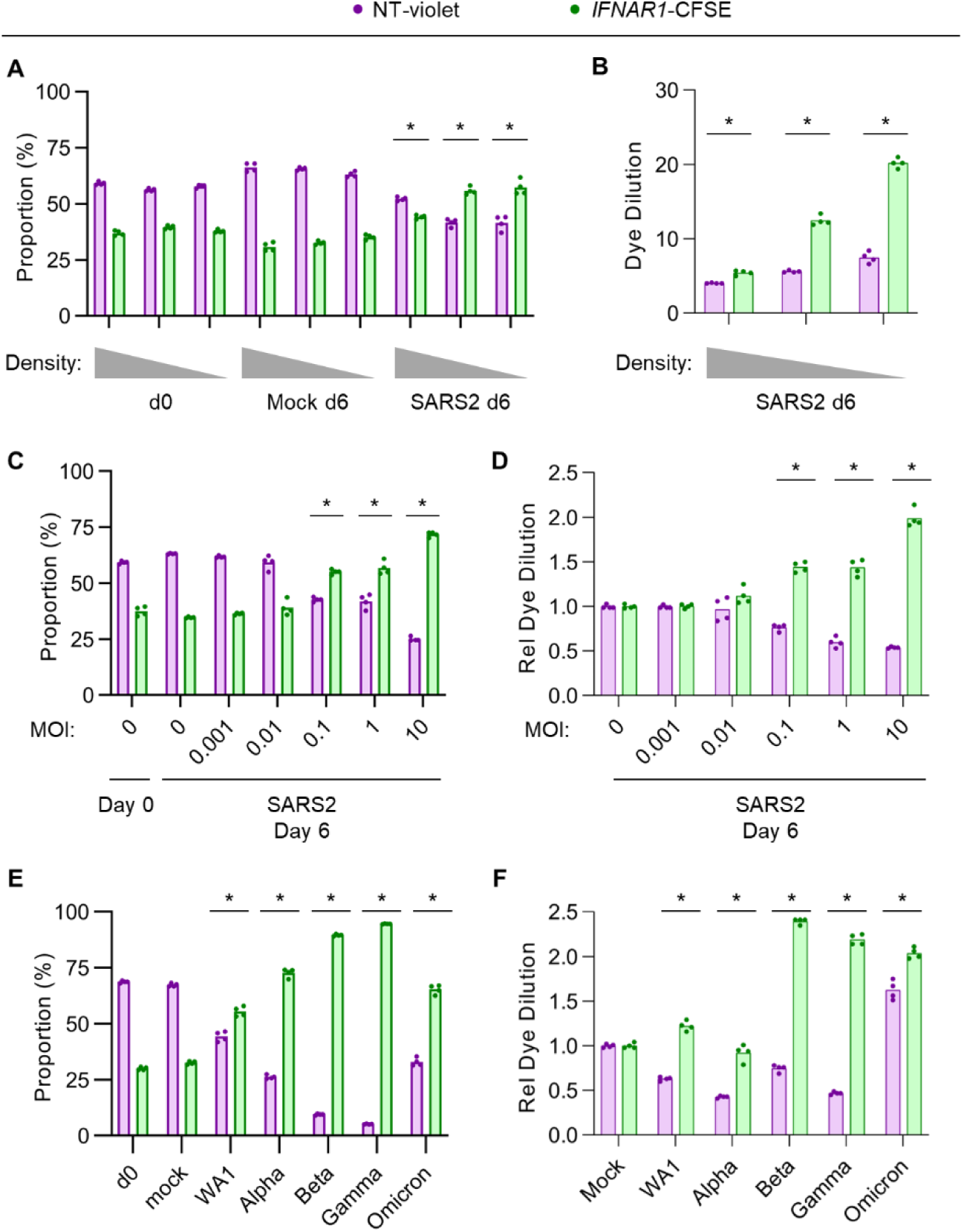
The competitive advantage conferred by IFNAR signaling disruption during SARS-CoV-2 infection is modulated by seeding density and viral MOI. Cocultures of nontargeting gRNA-treated, CellTrace-Violet-loaded Calu3 cells and *IFNAR1*-targeting gRNA-treated, CellTrace-CFSE-loaded cells were (A-B) seeded at a range of plating densities and infected with SARS-CoV-2 WA1 strain at MOI of 1 or (C-F) seeded at the same ∼20% plating density and infected with (C-D) SARS-CoV-2 WA1 strain at a range of MOI or (E-F) SARS-CoV-2 variants at MOI of 1. The relative proportion of each cell population was analyzed by flow cytometry at the indicated time points (A, C, E) and the dilution of either fluorescent dye was quantified by changes in mean fluorescence intensity, normalized to the dilution of either dye under mock-infected conditions. Individual data points represent independent infections and asterisks indicate p < 0.01 by Student’s t-test between SARS-CoV-2 and mock-infected conditions (A, C, E) or between nontargeting and IFNAR1-targeting gRNA (B, D, F).

### IFN-I signaling alone, in the absence of viral infection, is sufficient to induce an antiproliferative response in Calu-3 cells

The resistance of *IFNAR1*-disrupted cells to the delayed antiproliferative response during SARS-CoV-2 infection indicated that IFN-I signaling was necessary for this response. To test whether it was also sufficient, we next treated uninfected Calu-3 cells with exogenous β-IFN at a range of concentrations and measured the impact on cell viability and proliferation. We observed a clear dose-dependent decrease in viable cells upon β-IFN treatment (Fig 6A), with an estimated IC_50_ that was approximately 20-fold lower than the reported concentration of endogenous β-IFN for wild-type Calu-3 cells at the peak of SARS-CoV-2 infection^30^. A time course analysis revealed the onset of this effect between days 2 and 4 post-treatment (Fig 6B), similar to the kinetics observed during SARS-CoV-2 infection (Fig 4A). This effect was dependent on IFN-I signaling, as it was prevented by either treatment with baricitinib (Fig 6B) or by genetic ablation of *IFNAR1* (Fig 6C). Also consistent with our findings during SARS-CoV-2 infection (Fig 5A-B), the magnitude of the antiproliferative response was dependent on cell density, as Calu-3 cells plated at lower density exhibited a greater decrease in viable cells upon β-IFN treatment compared to cells plated at higher densities (Fig 6D). Likewise, the β-IFN-mediated decrease in viable cells was not associated with an increase in cell death, as measured by membrane exclusion of a viability dye (Fig 6E) or by LDH release to the extracellular environment (Fig 6F), but rather by a reduction in cellular proliferation, as measured by dilution of fluorescent dye over time (Fig 6G). These findings indicate that IFN-I signaling is both necessary and sufficient to induce an antiproliferative effect in Calu-3 cells during SARS-CoV-2 infection and suggest that this response may be generalizable to other pathologic states associated with persistent IFN-I signaling in the respiratory epithelium.

**Figure 6.**
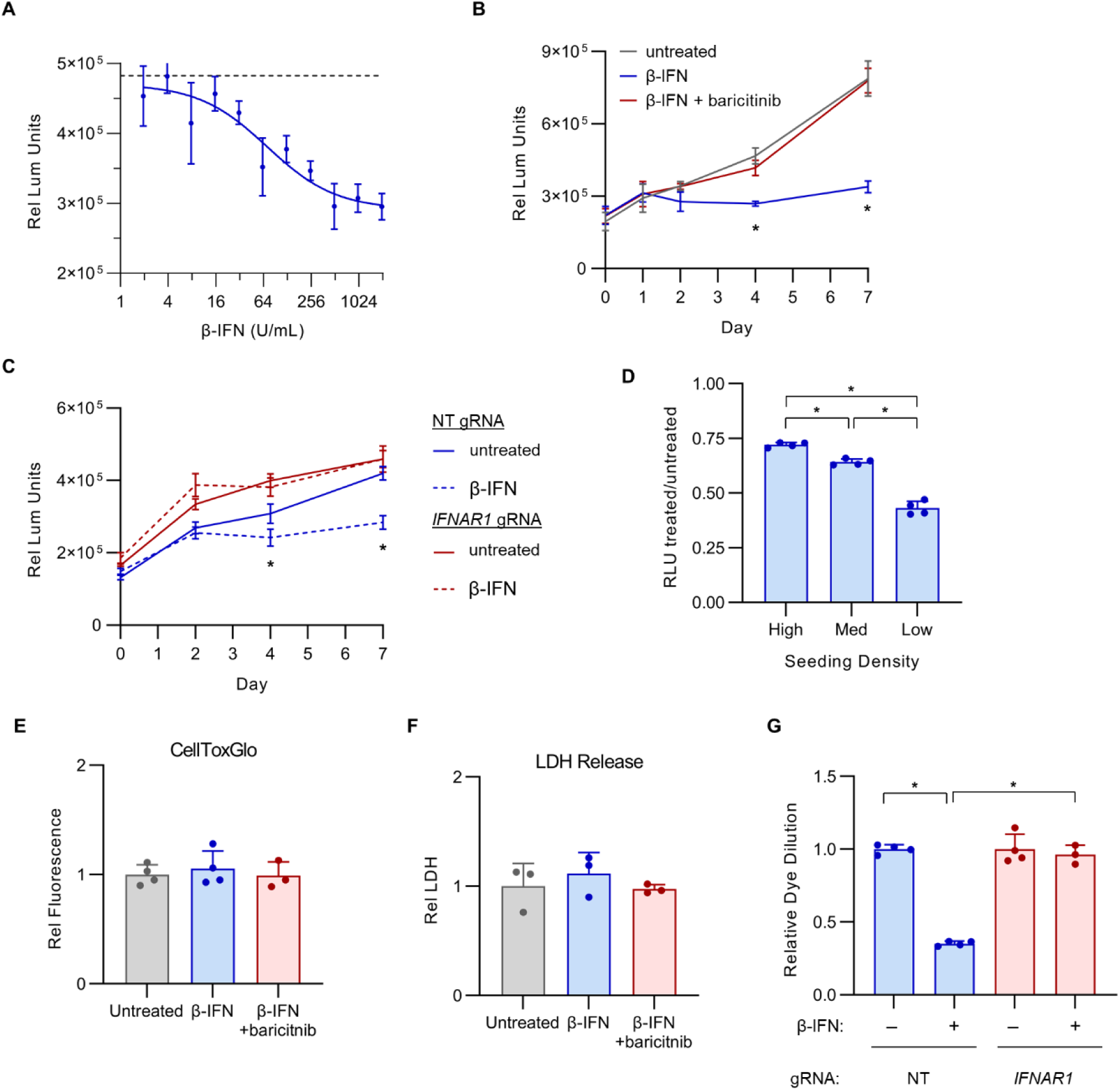
Exogenous IFN-I alone, in the absence of viral infection, is sufficient to induce an antiproliferative response in Calu-3 cells. (A) Calu-3 cells were incubated with a range of concentrations of exogenous β-IFN and the effect on cell quantities after 4 days were measured by CellTiter-Glo luminescence. (B) Calu-3 cells were incubated in the presence or absence of 1000 U/mL β-IFN with or without 1 µM baricitinib and analyzed by CellTiter-Glo luminescence at the indicated time points. (C) Calu-3 cells treated with a control nontargeting (NT) gRNA or a *IFNAR1*-targeting gRNA were incubated in the presence or absence of 1000 U/mL β-IFN and analyzed by CellTiter-Glo luminescence at the indicated time points. (D) Calu-3 cells were seeded at a range of densities (estimated confluence high = 80%, medium = 40%, low = 20%), incubated with or without 1000 U/mL β-IFN, and analyzed by CellTiter-Glo luminescence. The relative luminescence units of treated samples relative to untreated samples is plotted for each plating density. (E-F) Untreated and β-IFN-treated cells were analyzed for the presence of cell death after 4 days, as assayed by (E) cellular exclusion of the fluorescent viability dye CellToxGreen or (F) quantification of LDH released to the conditioned media. (G) The relative proliferation of untreated and β-IFN-treated Calu-3 cells were compared by flow cytometry detection of CellTrace-CFSE dilution over time. Error bars indicated standard deviations for technical replicates and asterisks indicate p < 0.001 by Student’s t-test.

## DISCUSSION

Because of its central importance in host defense against a wide variety of viral pathogens, the IFN-I response in COVID-19 has been the subject of extensive investigation since the onset of the pandemic. Several studies have found that pre-treatment of cultured cells with type I interferons before SARS-CoV-2 infection leads to a clear reduction in viral replication^39–43^ and this effect is mirrored by limited clinical trial data suggesting a benefit to patients for treatment with exogenous type I interferon prior to the onset of symptoms^44^. Human genetic studies have also suggested a link between interferon responses and COVID-19 outcomes, with one group finding 3.5% of patients with severe COVID-19 to harbor rare variants in interferon signaling genes (including *IFNAR1*, *IFNAR2*, *IRF3*, and *IRF7)*^45^, though this finding was not replicated in other studies^46, 47^. Autoantibody profiling of patients with severe COVID-19 also detected neutralizing antibodies against circulating interferons in 10.2% of patients^48^. Our finding that genetic or pharmacologic inhibition of IFN-I signaling leads to an increase in SARS-CoV-2 replication in Calu-3 cells is consistent with these studies demonstrating a protective effect for early IFN-I signaling during SARS-CoV-2 infection.

Despite the potential benefits of early IFN-I signaling during SARS-CoV-2 infection, several lines of evidence indicate that its persistent activation contributes to progression of lung disease in severe COVID-19. Longitudinal profiling of patients with COVID-19 has revealed a correlation between persistent IFN-I responses and greater severity of disease^49–54^. In mouse models of SARS-CoV-2 infection, genetic ablation of *Ifnar1*^55^ and pharmacologic inhibition of IFN-I production^56^ have each been shown to protect against lung pathology, while administration of exogenous IFN-I after the onset of symptoms in humans and in mice has been found to worsen disease^57–59^. A similar pathogenic effect for persistent IFN-I signaling has also been demonstrated in mouse models of SARS-CoV-1^60^ and MERS-CoV^61^ infection. Although specific IFN-I signaling inhibitors have not been tested in clinical trials, the Janus kinase inhibitors baricitinib and tofacitinib, which block IFN-I signaling (in addition to other pathways), have been shown to improve outcomes in severe COVID-19^14–16^. Together, these studies all support the overall pathogenic effect of persistent IFN-I signaling at advanced stages of SARS-CoV-2 infection, but do not identify the cell type or downstream pathways responsible for this effect.

Unexpectedly, our unbiased CRISPR screening approach revealed a cell autonomous and detrimental effect of persistent IFN-I signaling on Calu-3 cells during SARS-CoV-2 infection. The competitive advantage conferred by IFN-I signaling disruption in our screen might have been mediated by either a protection from SARS-CoV-2-induced cell death or an increase in cellular proliferation, and IFN-I signaling has been implicated in both processes in different cell types^62, 63^. We however found no influence for IFN-I signaling on Calu-3 cell death, with *IFNAR1*-disrupted cells showing equal sensitivity to SARS-CoV-2-induced death (Fig 4B) and exogenous IFN-I treatment causing no increase in cell death (Fig 6E-F). Instead, we found that the fitness advantage of *IFNAR1*-disrupted cells during SARS-COV-2 infection was mediated by an increase in cell proliferation between days 4 and 8 post-infection (Fig 4A, 4C), a finding that was mirrored by the antiproliferative response of uninfected cells to exogenous β-IFN treatment (Fig 6G).

Other high-throughput CRISPR screens for host factors affecting SARS-CoV-2 infection have not identified the same magnitude of protection as we observed for IFN-I signaling disruption. In some cases, this is likely due to the cell type tested. Vero E6 cells, for example, are deficient in the endogenous production of type I interferons^64^. Additionally, not all cell types exhibit an antiproliferative response to IFN-I signaling. Even for other screens of Calu-3 cells^27–29^, however, the observed effect for IFN-I signaling disruption has been variable (Fig S7). This is likely related to differences in screen design, as we identified multiple factors that influenced the relative advantage conferred by IFN-I signaling disruption. For example, Biering et al harvested cells at an earlier time point (day 5 post-infection rather than day 14 in our screen)^28^, which likely allowed for better detection of gene disruptions that affected cell death rather than the outgrowth of surviving cells. We likewise found that the magnitude of competitive advantage conferred by IFN-I signaling disruption was influenced by the plating density of the Calu-3 cells, the MOI of SARS-CoV-2 infection, and the SARS-CoV-2 variant tested. These discrepancies highlight how the technical nuances of screen design may bias toward the detection of host genes acting at different stages of viral infection.

Although our study is limited to the investigation of cell lines, the proliferation of respiratory epithelial cells is known to play a crucial role in recovery from acute lung injury *in vivo*, as progenitor cells (especially AT2 cells) generate a pool of cells that repopulate the injured alveolar lining^65^. The efficiency of this response is tightly linked to the clinical outcome of patients with acute respiratory distress syndrome^66–68^. Intriguingly, IFN-I signaling has been shown to negatively regulate AT2 proliferation in a mouse model of influenza^69^. It is unclear whether this interaction between IFN-I signaling and AT2 proliferation is unique to influenza or extends to other IFN-I-associated causes of acute lung injury. Our findings suggest IFN-I signaling alone is sufficient to cause an antiproliferative response in respiratory epithelial cells, supporting the potential generalizability of this effect. Our findings also suggest that the therapeutic benefit of Janus kinase inhibitors in severe COVID-19 may be due to their protection against an IFN-I-mediated antiproliferative response in respiratory epithelial cells. Future investigations will be necessary to establish the *in vivo* relevance of these findings and to explore the therapeutic potential of specifically targeting the IFN-I-mediated antiproliferative response during SARS-CoV-2 infection and other IFN-I-associated causes of acute lung injury and ARDS.

## Acknowledgments

This research was supported by the National Institutes of Health K08-HL148552 (BTE) and R01-GM139823 (AWT), the A. Alfred Taubman Medical Research Institute (BTE), the Michigan Institute for Clinical and Health Research (CM), the Marie-Slodowska Curie Global Fellowship GA-841247 (CM), and the University of Michigan Biological Scholars Program (CEW).

## Author Contributions

JBC, EJS, CM, CEW, and BTE conceived the project. EJS, SEG, CJW, AWT, JZS, CEW, and BTE designed the CRISPR library. EJS, CM, and BTE performed the CRISPR library synthesis and screening. JBC, EJS, CM, AAK, ASL, and BTE performed screen follow-up including generation and characterization of single gene-disrupted cell lines. JBC, CM, and AAK performed all BSL3 work. All authors contributed to data analysis and manuscript review. JBC and BTE wrote the manuscript with input from all authors.

## Declaration of interests

The authors have no relevant competing financial interests to declare.

## MATERIALS AND METHODS

### Cell lines and virus stocks

Calu-3 and HEK-293T cells were obtained from ATCC (Manassas, VA) and cultured in DMEM supplemented with 10% FBS, 10 U/mL penicillin, and 10 µg/mL streptomycin in a humidified 5% CO2 chamber at 37°C. Cell lines were periodically tested for mycoplasma contamination and identity confirmation by microsatellite genotyping. SARS-CoV-2 WA1 (isolate USA-WA1/2020, lineage A, catalogue NR-52281), Alpha (isolate USA/CA_CDC_5574/2020, lineage B.1.1.7, catalogue NR-54011), Beta (isolate hCoV-19/USA/MD-HP01542/2021, lineage B.1.351, catalogue NR-55282), Gamma (isolate hCoV-19/Japan/TY7-503/2021, lineage P.1, catalogue NR-54982), and Omicron (isolate hCoV-19/USA/MD-HP20874/2021, lineage B.1.1.529, catalogue NR-56461) variants were obtained from BEI Resources and propagated in Vero E6 cells. SARS-CoV-2 encoding nanoluciferase^38^ was kindly provided by Dr. R. Baric (University of North Carolina). Lack of genetic drift in viral stocks was confirmed by full-length sequencing as previously described^70^. Viral titers were determined by TCID50 assays in Vero E6 cells by microscopic scoring using the Reed-Muench method^71^. All experiments using SARS-CoV-2 were performed at the University of Michigan under Biosafety Level 3 protocols in compliance with containment procedures in laboratories approved for use by the University of Michigan Institutional Biosafety Committee (IBC) and Environment, Health and Safety (EHS).

### CRISPR screen design

The design and synthesis of the custom CRISPR library was previously described^31^. For each of 2 independent biologic replicates, a total of ∼50 million Calu-3 cells were transduced with the lentiviral CRISPR library at an MOI of ∼0.3. Puromycin was added at a concentration of 3 µg/mL at day 1 post-transduction and maintained until selection of control uninfected cells was complete. Cells were passaged as needed to maintain logarithmic phase growth with total cell number maintained above a minimum of 10 million cells at all stages of the screen. At 14 days post-transduction, a total of ∼40 million cells in 4 T175 flasks at approximately 50% density were infected with SARS-CoV-2 WA1 strain at MOI 1.0 in 10 mL infection media (DMEM with 2% FBS). Two days post-infection, with the incursion of visual cytopathic effect, 15 mL of maintenance media (DMEM with 10% FBS) was added. Cells were inspected every two days and medium was replaced at day 6 and 10 post-infection to allow enrichment of surviving cells. Cells were harvested at day 14 post-infection, genomic DNA was extracted with QIAamp DNA Mini Kit, and gRNA sequences were amplified and sequenced as previously described^72^.

### CRISPR screen analysis

FASTQ files were processed by PoolQ (Broad Institute; https://portals.broadinstitute.org/gpp/public/software/poolq) to map individual sequencing reads to reference gRNA sequences with deconvolution of sample identity by barcode. Cumulative distribution functions of gRNA representation were generated by plotting normalized read counts of each gRNA against its relative rank for a given barcode. Individual gRNA-level and aggregate gene-level enrichment analysis was performed using MAGeCK^35^. Q-Q plots were generated by plotting log-transformed observed p-values (calculated by MAGeCK gene-level analysis) against expected p-values (determined by the relative rank of each gene among the library). Genes were considered screen hits if they were identified with a MAGeCK-calculated false discovery rate < 0.05. Identified genes were filtered to remove those genes for which disruption influenced the fitness of uninfected cells, as reflected by a significant (FDR<5%) aggregate enrichment (log2FC>1) or depletion (log2FC<-1) of gRNAs prior to SARS-CoV-2 infection, relative to the plasmid pool. Genes targeted by depleted gRNAs were compared to the mean Gene Effect score in CRISPR screens of 1070 cell lines in the Broad Institute Cancer Dependency Map release 22Q1^36^. Enrichment of functional annotations among screen hits relative to all genes in the CRISPR library was performed using GOrilla with default setting of p-value < 10^-3^ for Gene Ontology Biologic Processes^73^; when multiple nodes were identified within the same hierarchy, the most significantly enriched term was selected for visualization. Gene network analysis of screen hits was performed using the STRING database^74^ with default settings and visualized using Cytoscape^75^ v3.9.1 with node borders weighted by -log(RRA score) for each gene in the CRISPR screen and edges weighted by gene-gene interaction score in the STRING database. Heatmaps were generated with GraphPad Prism v9.1.0 from published CRISPR screen data, using each gene’s aggregate gRNA log_2_-fold change or Z-score and the limits of the color spectrum assigned to values at the 99.9 and 0.1 percentiles within each screen.

### Generation of *IFNAR1*-disrupted and control nontargeting cells

CRISPR lentiviral stocks were generated by ligating a *IFNAR1*-targeting [GTACATTGTATAAAGACCAC] or control nontargeting sequence [GTTCATTTCCAAGTCCGCTG] into *BsmBI*-digested pLentiCRISPRv2^76^, cotransfecting each construct with psPAX2 and pVSVG into HEK293T cells, harvesting supernatants, and titering virus stocks as previously described^77^. Calu-3 cells were transduced in parallel with each lentiviral construct, treated with puromycin 3 µg/mL until no surviving cells remained among control non-transduced cells, and passaged to remain logarithmic phase growth. Phenotypic analysis was performed after a minimum of days 14 post-transduction to ensure adequate time for target site editing and turnover of residual protein.

### Analysis of cells with activation or inhibition of IFN-I signaling

Calu-3 cells were treated with exogenous β-IFN (R&D Systems, Minneapolis MN), baricitinib (Selleck Chemicals, Houston TX), or vehicle at the indicated time points and concentrations. Total RNA was isolated from cell lysates or the conditioned media using the RNeasy Plus Micro kit (Qiagen, Hilden Germany) or QIAamp Viral RNA kit (Qiagen), respectively, and converted to cDNA with the SuperScript III First-Strand Synthesis kit (Thermo Fisher, Waltham MA). Individual transcripts were amplified with primer pairs for *IFNAR1* (proprietary sequence, assay number Hs.PT.58.20048943, Integrated DNA Techologies, Coralville IA), *IFIT1* ([AAGCTTGAGCCTCCTTGGGTTCGT] and [TCAAAGTCAGCAGCCAGTCTCAGG]), *IFITM1* ([CCAAGGTCCACCGTGATTAAC] and [ACCAGTTCAAGAAGAGGGTGTT]), and SARS-CoV-2 RdRP (F2 primer IDT catalogue 10006860 and R1 primer IDT catalogue 1000688) using the Power SYBR Green PCR Master Mix (Thermo Fisher), and analyzed by QuantStudio 5 Real-Time PCR (Thermo Fisher). Quantification of individual transcripts was normalized against a panel of *ACTB*, *RPL37*, and *RL38* loading controls, as previously described^78^. Immunoblotting of RIPA lysates was performed as previously described^79^ with antibodies against IFNAR1 (Abcam, Cambridge UK, ab124764, 1:500), IFITM3 (Proteintech, Rosemont IL, 117141, 1:500), and β-actin (Santa Cruz Biotechnology, Dallas TX, sc-47778, 1:5000). The quantity of viable cells over time was monitored using CellTiter-Glo 2.0 (Promega, Madison WI) according to manufacturer’s instructions, with luminescence read on a VICTOR Nivo plate reader (PerkinElmer, Waltham MA). Cell death was monitored using LDH-Glo Cyotoxicity Assay (Promega), CellToxGreen (Promega), or flow cytometry of cells stained with LIVE/DEAD Fixable Far Red viability dye (Thermo Fisher).

### Competitive co-culture experiments

Control and *IFNAR1*-disrupted Calu-3 cells were detached using TrypLE Express, collected in DMEM with 10% FBS, centrifuged at 500g x 5 min, and the supernatant was aspirated. Cell pellets were then resuspended in 8 mL PBS pre-warmed at 37°C and CellTrace-Violet or CellTrace-CFSE added to the suspension to a final concentration of 5 µM. Cell suspensions were then incubated at 37°C for 20 min with periodic gentle agitation. Labeling was then quenched by the addition of 40 mL DMEM with 10% FBS at 4°C and cells centrifuged as above. The supernatant was aspirated and cell pellets resuspended in DMEM with 10% FBS. Cells were then pooled together and seeded into 6 well plates at a confluence of approximately 20% or as otherwise indicated. SARS-CoV-2 infections were performed 4 days later as described above. Media was replaced the day after seeding cells, the day prior to SARS-CoV-2 infection, and approximately every 3-4 days after infection. At the indicated time points, cells were detached, washed, stained with LIVE/DEAD Fixable Far Red viability dye (Thermo Fisher) for 15 minutes at 4°C, washed, fixed with 4% paraformaldehyde for 15 min at room temperature, and analyzed by flow cytometry. Where indicated, CountBright Absolute Counting Beads (Thermo Fisher) were spiked into samples at 1000 beads/µL immediately prior to flow cytometry. Flow cytometry was performed on a BioRad Ze5 instrument. Gates were established based on control populations of cells labeled with either individual dye and unlabeled cells that were cultured in isolation. Analysis of event gating frequency and mean fluorescence intensity was performed using FlowJo v10.8.1.

**Supplemental Table 1. Gene-level CRISPR screen results for modifiers of Calu-3 cell fitness during SARS-CoV-2 infection.** MAGeCK output for aggregate gene-level gRNA-level enrichment or depletion following SARS-CoV-2 infection relative to input cells. Negative log_2_-fold change and RRA scores indicate gRNA depletion in SARS-CoV-2-infected cells (gene disruption reduces cell fitness); positive values indicated gRNA enrichment in SARS-CoV-2-infected cells (gene increases cell fitness).

**Supplemental Table 2. Individual gRNA-level CRISPR screen results for modifiers of Calu-3 cell fitness during SARS-CoV-2 infection.** MAGeCK output for individual gRNA-level enrichment or depletion following SARS-CoV-2 infection relative to input cells. Negative log_2_-fold change and RRA scores indicate gRNA depletion in SARS-CoV-2-infected cells (gene disruption reduces cell fitness); positive values indicated gRNA enrichment in SARS-CoV-2-infected cells (gene increases cell fitness).

**Supplemental Table 3. Gene-level CRISPR screen results for modifiers of the fitness of uninfected Calu-3 cells.** MAGeCK output for aggregate gene-level gRNA-level enrichment or depletion in cells transduced with lentiviral pool for 2-3 weeks prior to onset of SARS-CoV-2 infection, relative to plasmid pool. Negative log_2_-fold change and RRA scores indicate gRNA depletion in SARS-CoV-2-infected cells (gene disruption reduces cell fitness); positive values indicated gRNA enrichment in SARS-CoV-2-infected cells (gene increases cell fitness).

**Supplemental Table 4. Individual gRNA-level CRISPR screen results for modifiers of the fitness of uninfected Calu-3 cells.** MAGeCK output for individual gRNA-level enrichment or depletion in cells transduced with lentiviral pool for 2-3 weeks prior to onset of SARS-CoV-2 infection, relative to plasmid pool. Negative log_2_-fold change and RRA scores indicate gRNA depletion in SARS-CoV-2-infected cells (gene disruption reduces cell fitness); positive values indicated gRNA enrichment in SARS-CoV-2-infected cells (gene increases cell fitness).

**Supplemental Table 5. Statistical analysis.** Source data and statistical analyses for the indicated figures.

---

**Figure S1.**
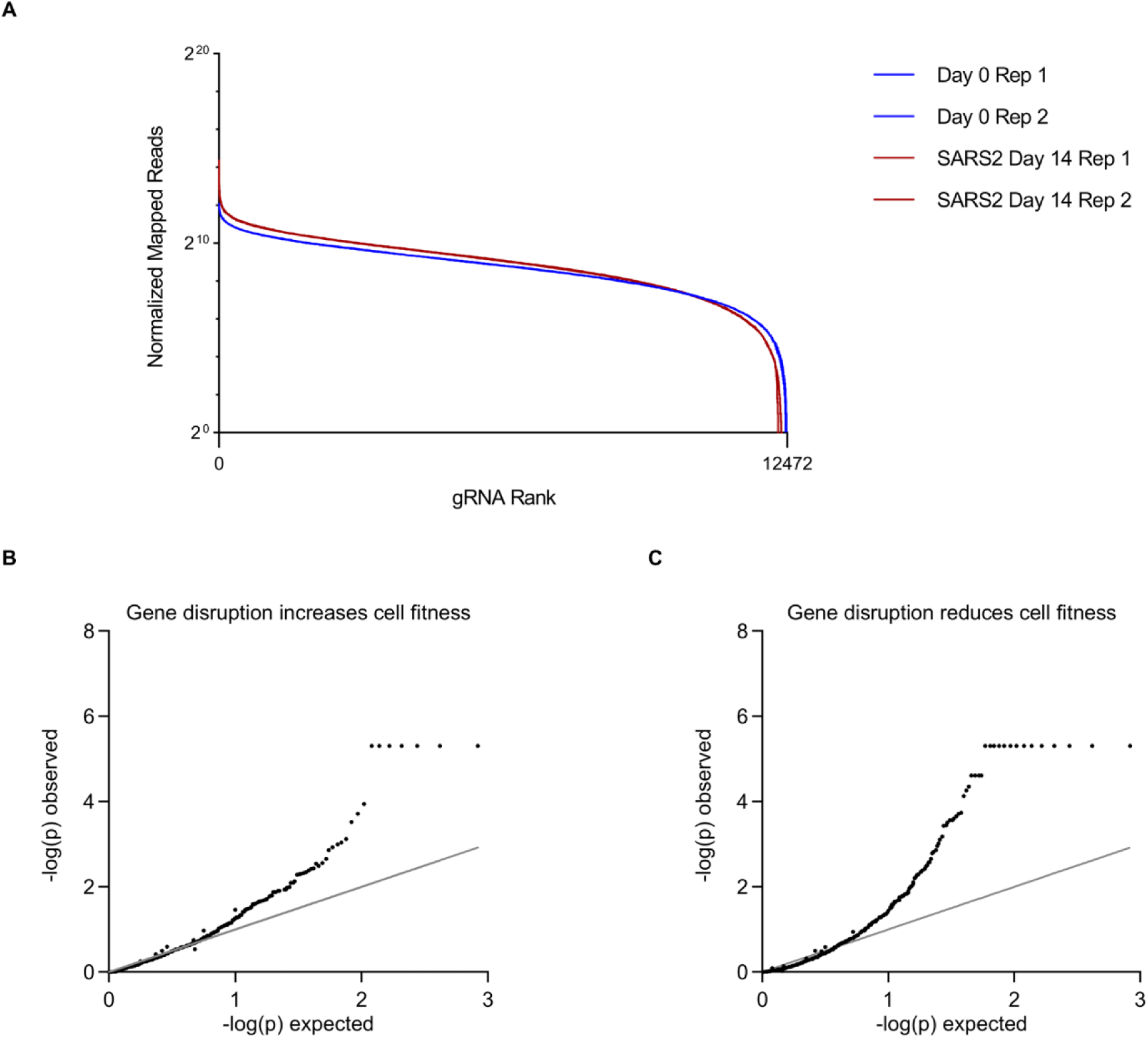
CRISPR screen analysis. (A) Cumulative distribution functions of normalized read counts for each gRNA before and after 14 days of SARS-CoV-2 infection for each of 2 independent biologic replicates. (B) Q-Q plots of observed versus expected –log(p) for gene perturbations associated with increased (B) or decreased (C) cell fitness during SARS-CoV-2 infection for every gene targeted by the CRISPR library. Observed p-values were calculated by MAGeCK gene-level analysis (Supplemental Table 1).

**Figure S2.**
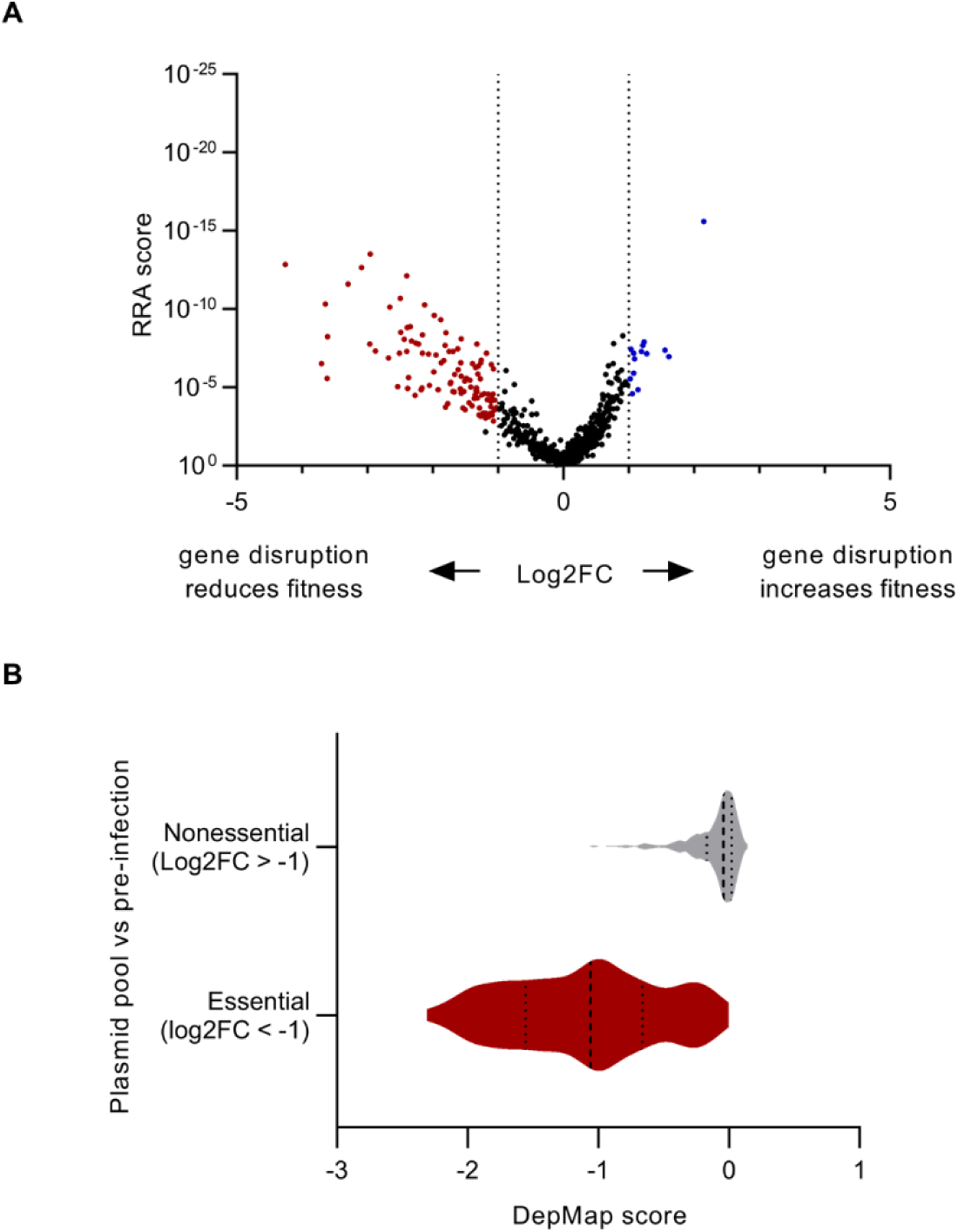
Filtering of genes with influence on Calu-3 cell fitness independently of SARS-Cov-2 infection. (A) Volcano plot of gRNA enrichment in day 0 pre-infected cells relative to the CRISPR library plasmid pool. Genes whose disruption conferred a significant (FDR<5%) increase (aggregate log2FC>1) or decrease (aggregate log2FC <-1) in cell fitness are highlighted in blue or red, respectively, and were filtered out of the CRISPR screen results for modifiers of cell fitness during SARS-CoV-2 infection. Source data is provided in Supplemental Tables 3 and 4. (B) Aggregate DepMap essentiality scores derived from CRISPR screens of 1070 cell lines for genes identified in this study as nonessential (L2FC>-1) or essential (L2FC<-1, FDR<5%) in Calu-3 cells prior to the onset of SARS-CoV-2 infection. DepMap scores of 0 indicate neutral effect on cell fitness while significant negative scores are consistent with a core essential function across cell lines.

**Figure S3.**
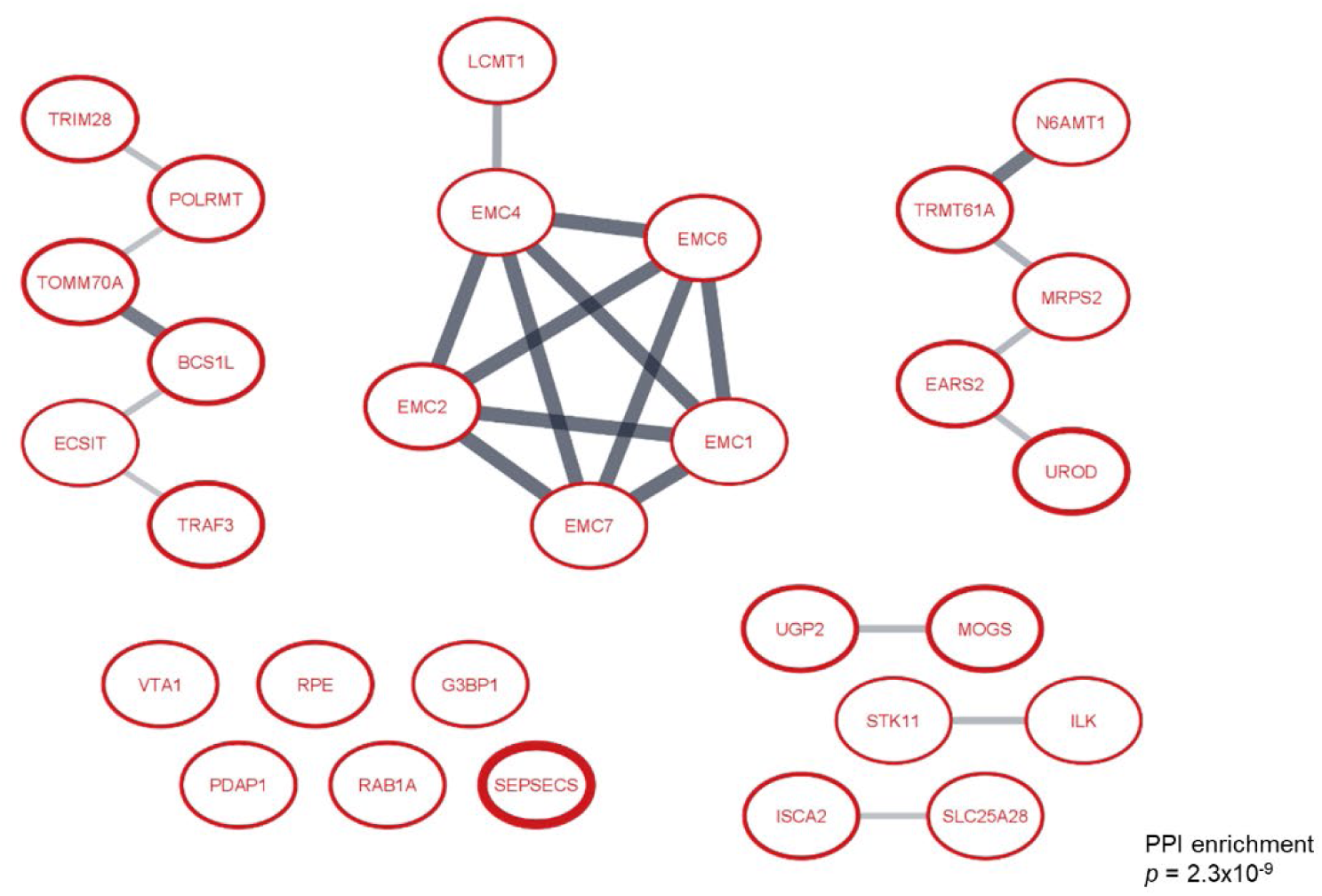
Established networks among genes whose disruption reduced the fitness of Calu-3 cells during SARS-CoV-2 infection. Borders of individual nodes are weighted by the - log(RRA score) in the screen, and lines connecting nodes are weighted by the strength of the protein-protein interaction within the STRING database. The significance of the number of protein-protein interactions relative to a randomly selected gene set was calculated by STRING.

**Figure S4.**
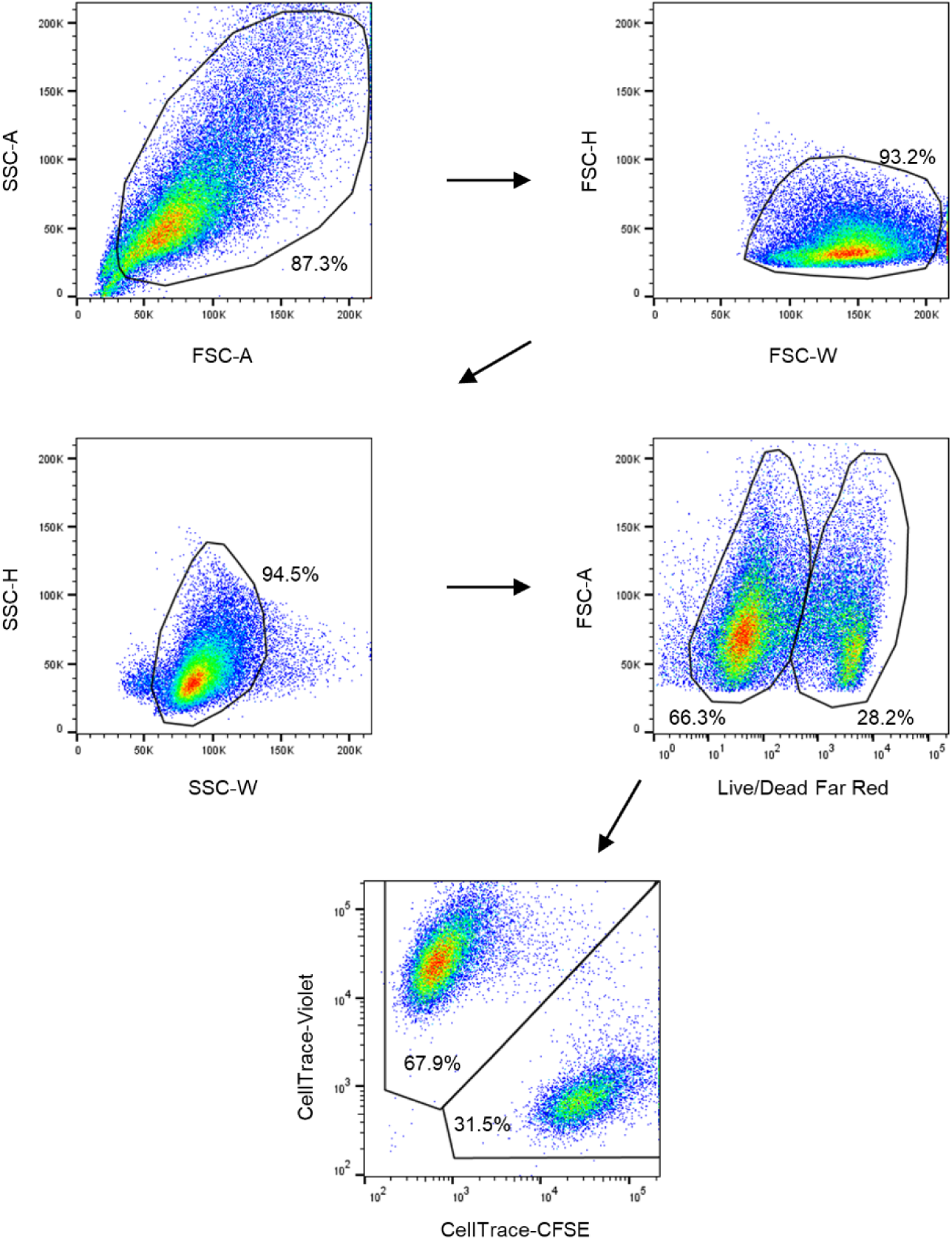
Representative flow cytometry plots. Gating strategy for quantifying relative proportions of cells labeled with CellTrace-CFSE and CellTrace-Violet, as indicated in Figures 2, 4, 5, 6, and S5.

**Figure S5.**
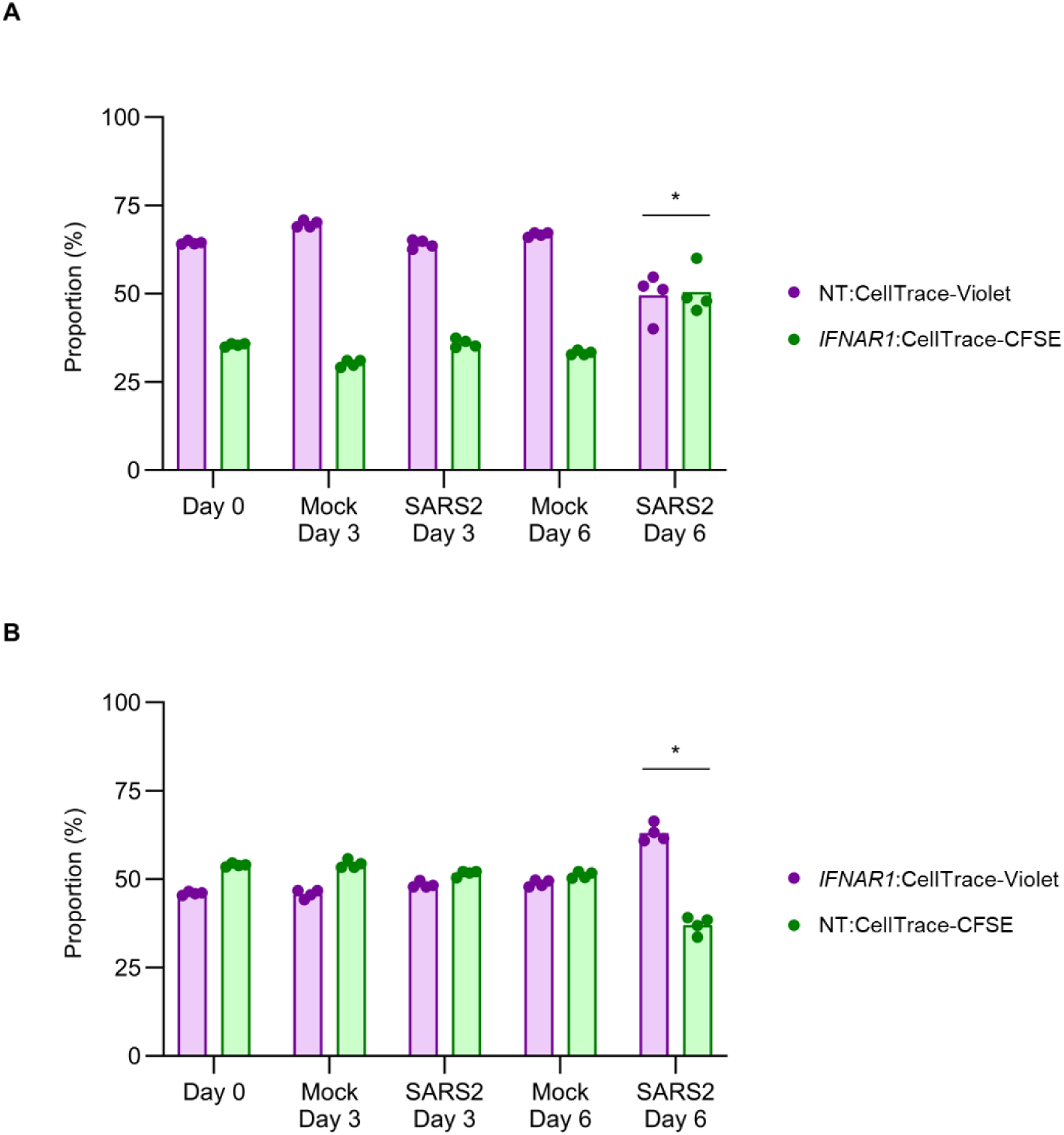
The competitive advantage of *IFNAR1*-disrupted cells in co-culture with control nontargeting gRNA-treated cells during SARS-CoV-2 infection is not mediated by the fluorescent tracker. Calu-3 cells treated with a nontargeting gRNA were labeled with either CellTrace-Violet (A) or CellTrace-CFSE (B) and seeded in co-culture with cells treated with a *IFNAR1*-targeting gRNA and labeled with the other fluorescent tracker, CellTrace-CFSE (A) or CellTrace-Violet (B). The relative abundance of each cell population was analyzed by flow cytometry at the indicated time points under conditions of mock or SARS-CoV-2 infection. Data points represent independent infections and asterisks indicate p < 0.05 for the relative proportions of each population between the indicated groups by Student’s t-test.

**Figure S6.**
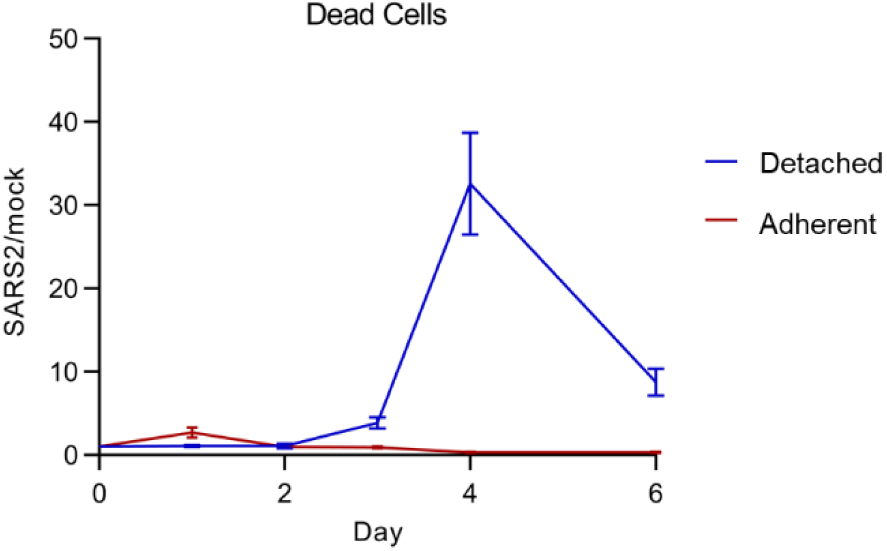
Quantification of adherent and detached dead cells over the course of mock or SARS-CoV-2 infection. The ratio of dead cells associated with either the cell monolayer or detached in the conditioned media is plotted for SARS-CoV-2 infection relative to mock infection conditions at the indicated time points. Error bars represent standard deviation of 4 independent infections.

**Figure S7.**
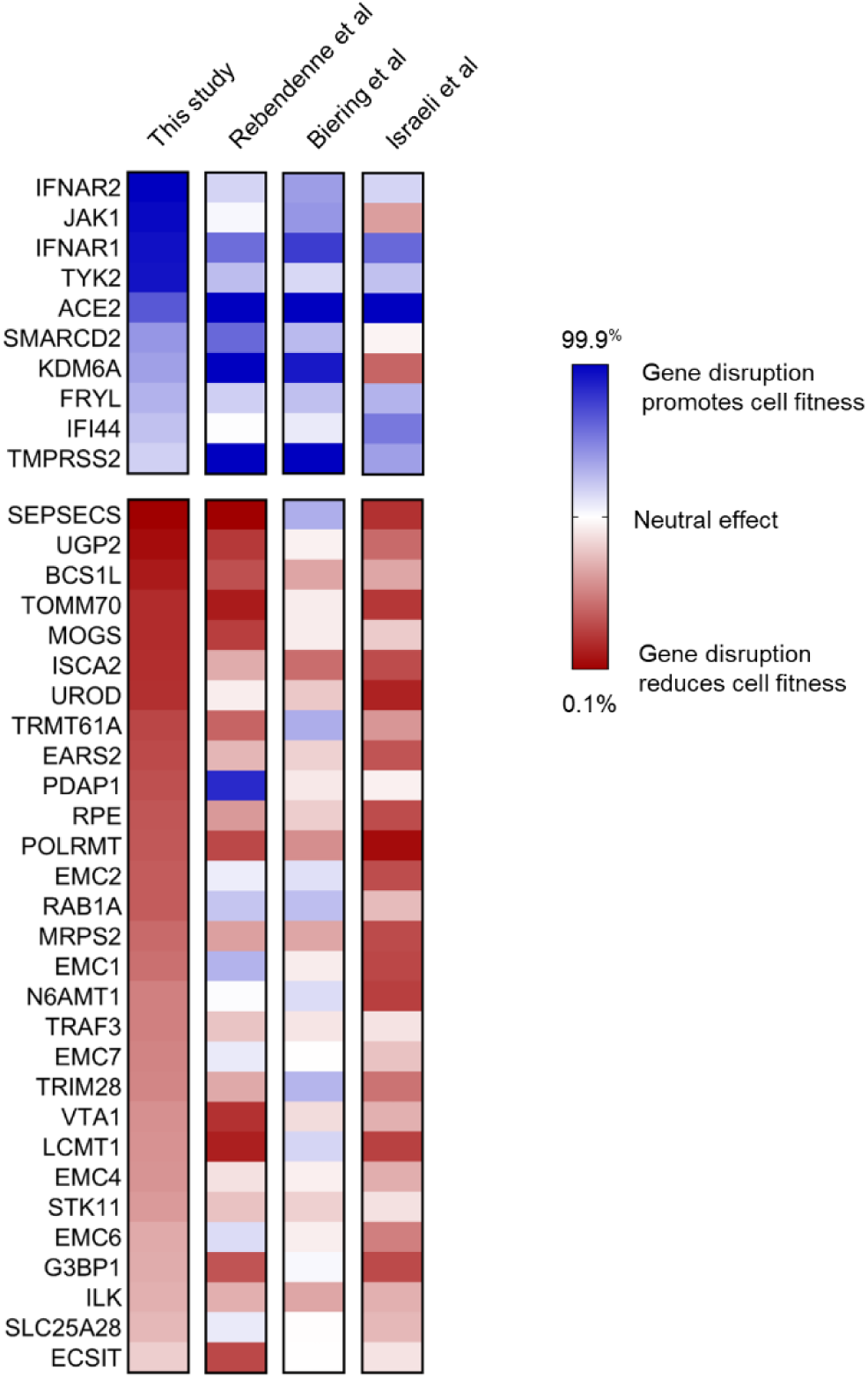
Comparative analysis of genes identified in this study with other published CRISPR screens of Calu-3 cell fitness during SARS-CoV-2 infection. Heat map of aggregate gRNA enrichment (blue) or depletion (red) for the indicated genes in each CRISPR screen. Values from other published studies were normalized with the end of each color spectrum representing the 99.9 and 0.1 percentiles among all genes queried in the screen, placed in rank order by log2FC or Z-score.

## Notes

### Competing Interest Statement

The authors have declared no competing interest.

